# Co-expression patterns of microglia markers Iba1, TMEM119 and P2RY12 in Alzheimer’s disease

**DOI:** 10.1101/2021.05.31.446375

**Authors:** Boyd Kenkhuis, Antonios Somarakis, Lynn RT Kleindouwel, Willeke MC van Roon-Mom, Thomas Höllt, Louise van der Weerd

## Abstract

Microglia have been identified as key players in Alzheimer’s disease pathogenesis, and other neurodegenerative diseases. Iba1, and more specifically TMEM119 and P2RY12 are gaining ground as presumedly more specific microglia markers, but comprehensive characterization of the expression of these three markers individually as well as combined is currently missing. Here we used a multispectral immunofluorescence dataset, in which over seventy thousand microglia from both aged controls and Alzheimer patients have been analysed for expression of Iba1, TMEM119 and P2RY12 on a single-cell level. For all markers, we studied the overlap and differences in expression patterns and the effect of proximity to β-amyloid plaques. We found no difference in absolute microglia numbers between control and Alzheimer subjects, but the prevalence of specific combinations of markers (phenotypes) differed greatly. In controls, the majority of microglia expressed all three markers. In Alzheimer patients, a significant loss of TMEM119^+^-phenotypes was observed, independent of the presence of β-amyloid plaques in its proximity. Contrary, phenotypes showing loss of P2RY12, but consistent Iba1 expression were increasingly prevalent around β-amyloid plaques. No morphological features were conclusively associated with loss or gain of any of the markers or any of the identified phenotypes. All in all, none of the three markers were expressed by all microglia, nor can be wholly regarded as a pan- or homeostatic marker, and preferential phenotypes were observed depending on the surrounding pathological or homeostatic environment. This work could help select and interpret microglia markers in previous and future studies.

## 1. Introduction

Microglia are the resident myeloid cells of the central nervous system, crucial for both development and homeostasis of the brain. With their highly motile processes, they continuously survey their micro-environment (1), enabling them to not only act as first in line immune defense, but also help regulate brain activity (2). Microglia are also implicated to play an important role in several neurodegenerative diseases, including Alzheimer’s disease (AD) (3). Genome wide association studies have identified many risk genes for AD, with the majority primarily or even exclusively expressed in microglia, indicative of a causal or augmentary role for microglia in disease pathogenesis(4). In line with this, transcriptomic studies have found microglia to show the most substantial changes in response to pathology in murine and human studies (5–8). These studies highlight the importance of well-characterized tools to study microglia in both homeostasis and disease.

One of the most widely used techniques to study microglia is immunohistochemistry (IHC). For a long time, microglial researchers depended on classical myeloid cell markers, of which the most commonly used ones are ionized calcium binding adaptor molecule (Iba)1, MHCII cell surface receptor HLA-DR, CD11b and CD68. However, these markers do not allow distinguishing microglia from infiltrating macrophages. The importance of this distinction has become increasingly clear, as microglia arise from embryonic yolk sac precursors during early embryonic development, which makes them ontogenically and transcriptionally distinct from blood-associated myeloid cells such as macrophages, that infiltrate the human CNS under pathological conditions (9). Recently, researchers identified new genes that are almost exclusively expressed by microglia in CNS (10). The two most prominent examples of such genes are *TMEM119* and *P2RY12*, for which quickly novel antibodies were developed.

Transmembrane protein 119 (TMEM119), was characterized to be a specific microglia marker by Bennet et al in 2016, although its function in microglia remains unknown (11). It is considered to be a marker expressed by microglia under homeostatic conditions based on transcriptomic findings (5,6,12), but also *in vitro* mRNA levels are reduced when microglia are activated (11). However, contrary to these results, increased mRNA levels have been found in AD brains (13). P2RY12 on the other hand, is a purinergic receptor that responds to ADP/ATP to increase cell migration. It was initially identified on platelets, and plays an important role in mediating platelet activation and blood clotting (14). P2RY12 was found to be exclusively expressed by microglia in the murine CNS (10), and consistently expressed by human microglia throughout development(15). Its expression is decreased under pathological conditions present in AD (15), which Walker *et al*. later attributed to a specific lack of P2RY12-positive cells surrounding mature amyloid beta (Aβ) plaques (16). However, P2RY12 expression was also found on microglia positive for CD68 and HLA-DR, which are considered markers of activation(16). These findings suggest that TMEM119 and P2RY12 expression could encompass a wider range of microglia phenotypes than the originally hypothesized homeostatic subset.

While immunohistochemical evaluation of microglia using TMEM119 and P2RY12 is becoming more popular considering their microglia specificity, comprehensive data characterizing the expression in post-mortem AD brain tissue, as exists for the Iba1, CD68 and HLA-DR, is missing (17). Additionally, it is unknown what the overlap is between TMEM19 and P2RY12, and the classical marker Iba1. Therefore, in this study we aimed to investigate the expression patterns of Iba1, TMEM119 and P2RY12 in AD and age-matched control tissue and more importantly the overlap and differences in expression patterns. To do so, we used a previously acquired multispectral immunofluorescence dataset published by Kenkhuis *et al*. (18). On this dataset we performed a novel analysis using information about the expression of Iba1, TMEM119 and P2RY12 on a single-cell level and combined this with the spatial relationship with respect to Aβ plaques. We found a significant overall decrease of TMEM119 positive microglia in AD patients, but a strong loss of P2RY12 positivity and increase of Iba1 positivity specifically in Aβ-plaque surrounding microglia.

Our findings highlight the differences in expression between these markers in response to pathology, and will help choose the appropriate microglia marker for future research.

## 2. Methods

### 2.1 Study design

This study was performed using the multispectral immunofluorescence dataset published by Kenkhuis *et al*. on brain autopsy tissue of the middle temporal gyrus of 12 AD patients and 9 age-matched non-neurological controls (18). Summary of groups statistics can be found in table 1 or individual patient-characteristics can be found in Kenkhuis *et al*.. One section per subject was stained for a combination of markers for nuclei (DAPI) microglia (P2RY12, TMEM119 and Iba1) and Aβ-plaques.

**Table 1.** Summary of patient statistics. Difference in age of death and post-mortem interval was tested with a two-tailed Student’s independent t-test. Braak stage and Thal phase with a Mann

|  | Control (n = 9) | Alzheimer's disease (n = 12) | P-value |
| --- | --- | --- | --- |
| Age of onset, mean y (SD) | n.a | 61.9 (16.9) |  |
| Age of Death, mean y (SD) | 82.3 (8.3) | 74.8 (14.4) | 0.1754 |
| Male / female | 4/5 | 3/9 | 0.3972 |
| Braak stage, median (range) | 1/2 (0 - 3) | 6 (4 - 6) | < 0.0001 |
| Thal Phase, median (range) | 1 (0 - 2/3) | 5 (3/4 - 5) | < 0.0001 |
| Post-mortem interval, mean (SD) | 6:56 (1:31) | 4:49 (0:53) | 0.001 |
| APOE 2/3 - 3/3 - 3/4 - 4/4 | 0 - 3 - 1 - 0 | 1 - 5 - 4 - 2 |  |

Whitney-U test and male / female distribution with a Fisher's exact test.

Subsequently all microglia were segmented and raw intensity fluorescence values for all intracellular markers were extracted. Additionally, all parenchymal Aβ-plaques were segmented and the distance of each identified microglia to a parenchymal plaque was reported. Finally, each identified microglia was also assigned a number, which can be traced back to the original image to evaluate marker expression and morphology manually.

### 2.2 Multispectral immunofluorescence

In short, 5-μm-thick sections from Formalin fixed paraffin embedded (FFPE) tissue were deparaffinized with xylene and rehydrated using a gradient of 100%-50% alcohol. Endogenous peroxidases were blocked for 20 min in 0.3% H_2_O_2_/methanol and antigen-retrieval was performed by boiling the slides in pre-heated citrate (10 mM, pH = 6.0) for 10 min. Subsequently slides were blocked with blocking buffer (0.1% BSA/PBS + 0.05% Tween) for 30 min. Slides were then incubated sequentially with TMEM119 (1:250, Sigma Aldrich; Overnight) and P2RY12 (1:2500, Sigma Aldrich; 2 hour), each of which were amplified firstly with Poly-HRP secondary antibody and secondly with an Opal tertiary antibody, before continuing to the next antibody. Subsequently, slides were incubated overnight with a cocktail of Ferritin Light-chain (1:100, Abcam), Aβ (1:250, Biolegend) and Iba1 (1:20, Millipore) antibodies, diluted in blocking buffer. The next day, slides were incubated with a secondary antibodies for G-a-rIgG A594, G-a-mIgG2b A647 and G-a-mIgG1 CF680 (1:200, ThermoFisher), diluted in 0.1% BSA/BPS. Finally, the slides were washed and incubated with 0.1 μg/mL DAPI (Sigma Aldrich) for 5 min, after which they were mounted with 30 uL Prolong diamond (ThermoFisher). A more extensive step-by-step protocol for the multispectral immunofluorescence panel can be found in Kenkhuis *et al*. (18).

### 2.3 Data processing

Microglia were identified and segmented using a previously published inhouse cell segmentation pipeline (18). Raw fluorescence intensity values were normalized imposing Z-score transformation. To make our multispectral data comparable to traditional IHC studies, in which cells are usually scored as positive or negative for a certain marker, we aimed to set a threshold for positivity of an individual marker in our multispectral immunofluorescence data (Fig. 1A,B). We manually scored a subset of segmented microglia (n = 266) for cell-positivity of P2RY12, TMEM119 and Iba1, using a cell-positivity scoring scale (1: no signal; 2: sub-threshold positive, 3: positive signal, 4: very positive signal). Scoring was performed by two independent raters (BK and LRTK), after which a consensus score was reached (Supplementary Fig. 1A). By using the visually scored cells as ground truth values, we determined the optimal threshold values by calculating which values yielded the greatest percentage of true positive and true negative cells (Supplementary Fig. 1B). The calculated threshold values lead to true positive and true negative percentages of respectively 91% and 90% for P2RY12, 83% and 84% for TMEM119 and 86% and 84% for Iba1. Based on these calculated threshold values, all microglia could be assessed for positivity of the individual markers, and clusters could be created based on the combination of expression of the different markers (P2RY12^-/+^TMEM119^-/+^Iba1^-/+^).

**Fig 1.**
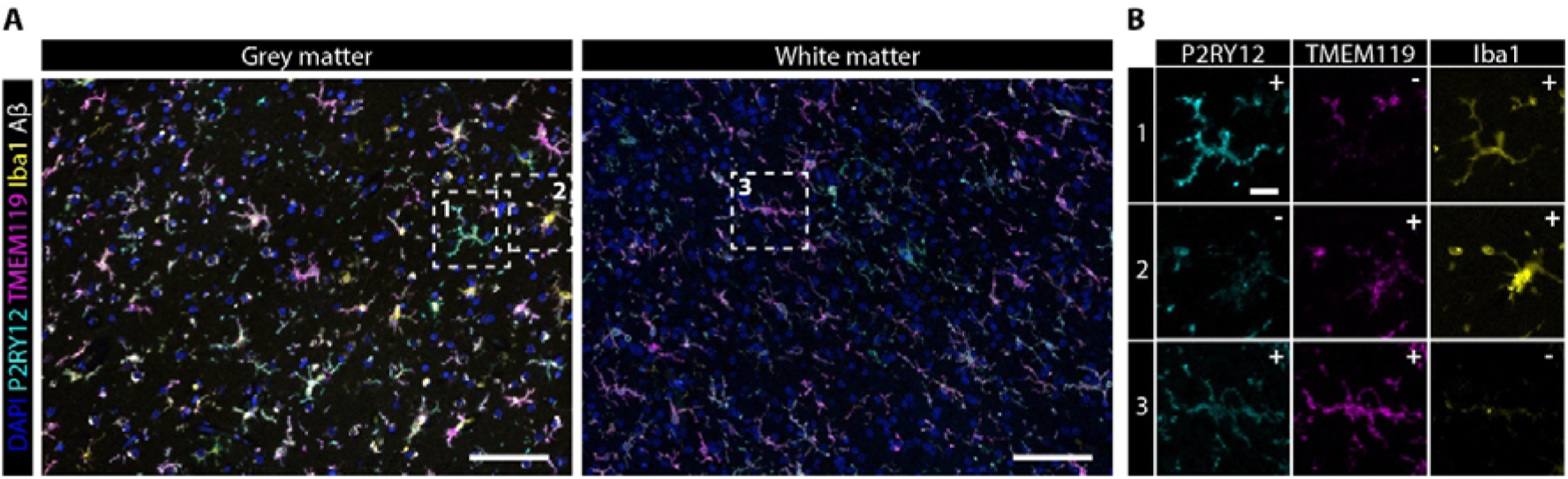
Co-expression of P2RY12, TMEM119 and Iba1 in multispectral immunofluorescence data. **A** Example of multispectral data of grey- and white matter of a control subject. **B** Zooms of individual microglia cells with manual evaluation for positive expression of different markers. Scale bar, 100 µm. Scale bar zoom, 20 µm.

### 2.4 Statistical analysis

All variables were inspected for gaussian distribution of the data. Normally distributed data are represented by mean and standard deviation, whereas not normally distributed data are represented with the median and corresponding interquartile range, which is stated in the figure legend.

Comparison of two continuous variables was performed using a two-tailed Student’s independent t-test (normally distributed) or a Mann-Whitney U test (not normally distributed). Paired spatial data was analysed using a Repeated Measures ANOVA with Geisser-Greenhouse correction. Bonferroni post-hoc analysis was performed on individual analyses, and a significance level of *P* < 0.05 was applied. All statistical tests were performed using GraphPad Prism (Version 8.00, La Jolla, San Diego, CA, USA).

## 3. Results

### 3.1 Differing prevalence of phenotypes in control and AD patients

In total, 76.512 myeloid cells (n = 1419-6500 per subject) were identified in 21 subjects, which were positive for one or more of the microglia markers. No significant differences were observed in the total number of identified microglia between controls and AD patients, neither in the grey or white matter (Figs. 2A,B). Although the total number of microglia did not differ between control and AD tissue, we did observe differences in the expression of the individual markers between controls and AD patients. The percentage of TMEM119^+^-microglia was significantly reduced in AD, both in the grey and in the white matter (Figs. 2C,D). For Iba1 and P2RY12 we did not observe significant differences, although P2RY12^+^-microglia appeared to be less present in the grey matter AD patients (Fig. 2C). We also compared the percentage of positive microglia for each individual marker between grey and white matter, but did not find significant differences between the grey- and white matter cell populations. When looking at the raw fluorescent intensity of the different markers in positive cells, which is considered to be correlated with protein quantity, we found the mean intensity for P2RY12 and TMEM119 to be decreased in AD (Fig. 2E). We also studied whether combinations of these different markers (‘phenotypes’) were more or less present in AD (Fig. 2F). Interestingly, even though the overall percentage of Iba1^+^-microglia was not increased in AD patients, microglia showing solely Iba1 expression without TMEM119 and P2RY12 (P4), were significantly more present in AD patients in both grey and white matter (Figs 1G,H,I). Conversely, P2RY12^+^TMEM119^+^Iba1^+^-microglia (P7) were significantly less prevalent in AD in the grey matter (Fig 1G,H). In the white matter specifically, P2RY12^+^-(P1) and P2RY12^+^Iba1^+^-microglia (P5) were significantly more present in AD (Fig 1I). Prevalence of phenotypes of all individual subjects that did not differ significantly, can be found in supplementary Fig. 2. These findings also show how the observed differences on single-marker level can be attributed to the combined differences in prevalence of the phenotypes. For both Iba1 and P2RY12, the increases observed in P1, P4 and P5 in AD are likely compensated by the decrease of P7 (Fig. 2H,I), resulting in no significant difference on single-marker level (Fig. 2C,D). For TMEM119 a significant decrease on single-marker levels was found, which can be attributed to the consistent findings of TMEM119^-^-phenotypes to show increased prevalence (P1, P4, P5) and TMEM119^+^-phenotypes (P7) to show decreased prevalence in AD (Fig. 2H,I).

**Fig 2.**
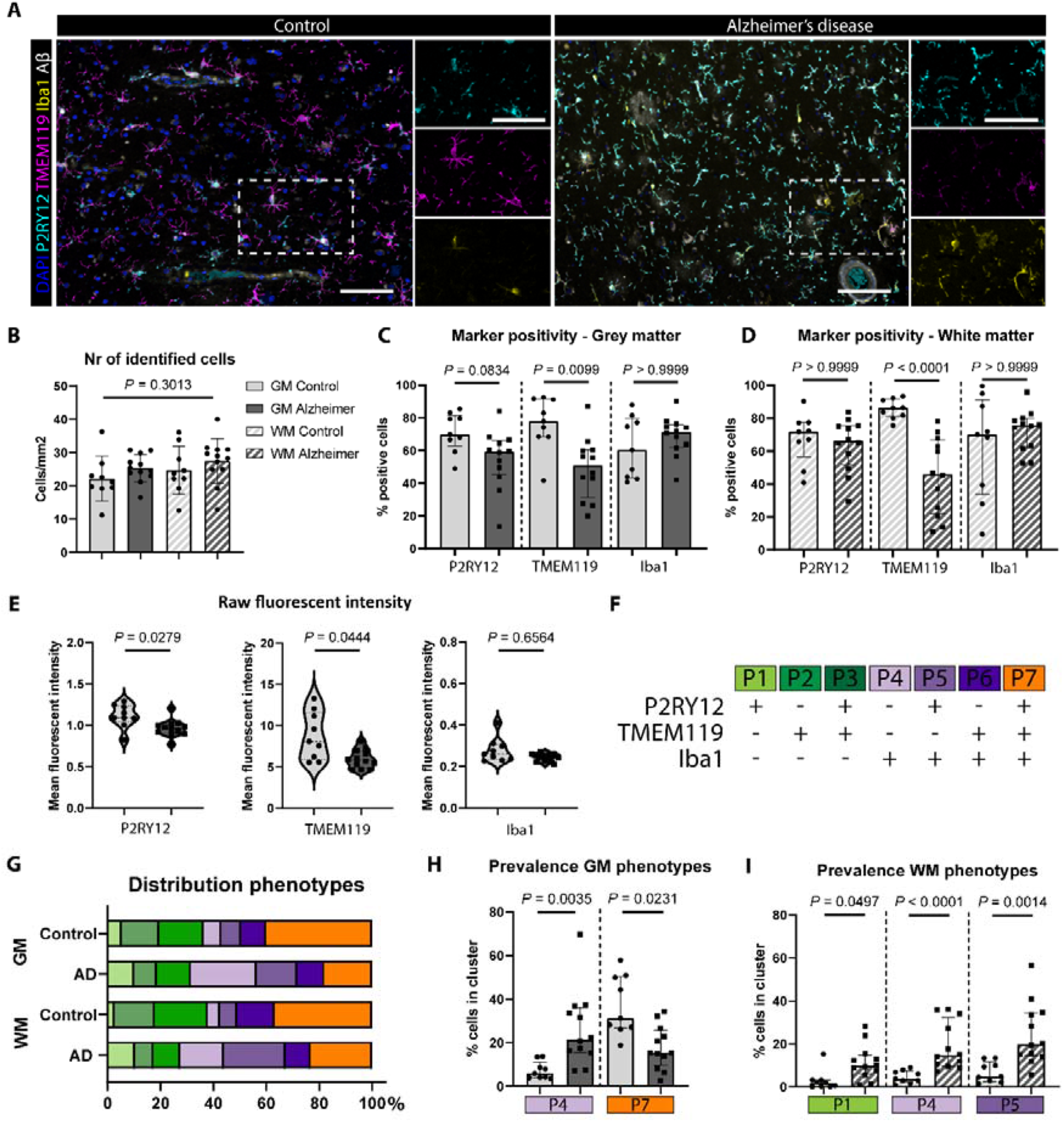
Consistent loss of TMEM119^+^- and variable increase of P2RY12^+^/Iba1^+^ phenotypes in Alzheimer brains. **A** Example image of control and AD patient with zooms showing individual markers **B** Number of identified cells in the grey- and white matter of controls (n = 9) and AD patients (n = 12) (Mean, One-way Anova). Percentage of cells positive for the individual markers in grey matter (**C**) and white matter (**D**) (Median, Mann Whitney U test). **E** Mean fluorescent intensity of positive cells for P2RY12, TMEM119 and Iba1 (Median, Mann Whitney U-test). **F** Created phenotypes (P) based on the combination of microglia markers present. **G** Distribution of identified phenotypes (P1–P7) in control and AD patients in the grey- and white matter. Prevalence of specific phenotypes differed significantly for in both the grey matter (GM) (**H)** and white matter (WM) (**I**) (Median, Mann-Whitney U test). *P*-values were corrected for multiple testing using Bonferroni within one graph (C, D, E, H and I). Scale bars, 100 µm.

### 3.2 P2RY12-expression but not TMEM119 is negatively associated with Aβ-plaques

After analyzing the differences in expression of the microglia markers between control and AD patients, we investigated whether the presence of Aβ-plaques changed the expression patterns of our three microglia markers. First, we calculated the distance of all microglia with respect to Aβ-plaques and divided them into three groups: infiltrating (0 µm), surrounding (0 – 20 µm) and no-plaque (>20 µm) (Fig. 3A). Again, we first evaluated expression of the single microglial markers. Whereas the decrease in TMEM119^+^-microglia in AD was again confirmed, no association with proximity to Aβ-plaques was observed (Fig. 3B). Conversely, there was a significant reduction in the number of P2RY12^+^-microglia infiltrating Aβ-plaques in AD patients and an increase in the cells positive for Iba1 in the plaque-surrounding microglia and even more so in the plaque-infiltrating microglia in both controls and AD patients (Fig. 3B). Next we investigated the effect of proximity to Aβ-plaques on the previously defined phenotypes. In controls, there was a decrease of P1-P3 and increase of P4 and P7 for microglia getting closer to Aβ-plaques (Fig. 3C). In AD patients, also a decrease in P1-P3 was observed. However, contrary to the controls, this was due to an increase of P4 and P6-microglia (Fig. 3C). This clearly indicates that the majority of Aβ-plaque infiltrating microglia consistently express Iba1, but variably show loss of TMEM119 and P2RY12. Subsequently we further looked into the observed trends in the individual subjects. None of the differences found in the control patients were found to be significant (Fig. 3D-J), which can be partially explained by the fact that fewer Aβ-plaques are identified in control patients, and therefore the accuracy of the phenotype distribution is much lower. In AD patients, consistent patterns are observed within individuals. For the Iba1^-^-phenotypes (P1-P3), P2RY12^+^-microglia (P1) showed the most significant decrease in prevalence when getting closer to Aβ-plaques (Fig. 3D-F). For the Iba1^+^-phenotypes (P5-P7), both Iba1^+^- (P4) and TMEM119^+^Iba1^+^-microglia (P6) showed significantly increased prevalence in microglia infiltrating Aβ-plaques (Fig. 3G,I). Two examples of such phenotypes infiltrating microglia are shown in Figs. 3K and 3L respectively. Notably, even though overall prevalence of Iba1 was increased for infiltrating microglia, P2RY12^+^Iba1^+^-microglia (P5) were significantly decreased in prevalence in Aβ-plaque infiltrating microglia. All in all, the presence of Aβ-plaques had the strongest effect on P2RY12-positivity, which was consistently decreased in plaque-infiltrating microglia (P1, P3, P5, P7). Most Iba1^+^-phenotypes were increased (P4, P6), though not consistently for all Iba1^+^-phenotypes (P5). No clear effect of the presence of Aβ-plaques was observed on the percentage of TMEM119^+^-microglia.

**Fig 3.**
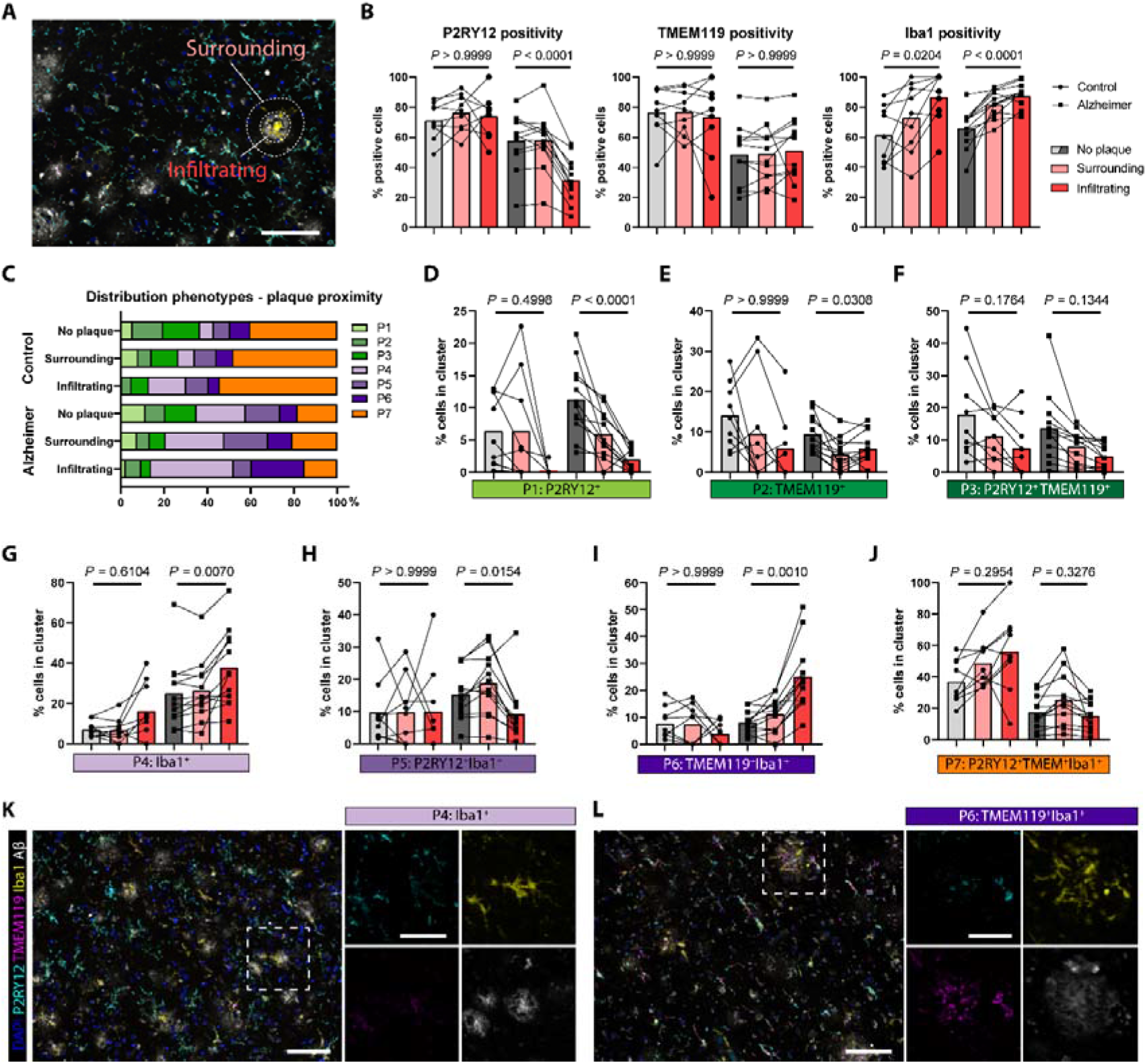
Preferential Aβ-plaque infiltration of Iba1^+^ and P2RY12^-^-phenotypes in Alzheimer patients. **A** Example of how microglia are spatially subdivided into surrounding (Pink; 0 < x < 20 um), infiltrating (Red; 0 um) or no plaque (> 20 um). **B** Prevalence of individual microglia markers in subdivided spatial groups of controls (n = 9) and AD patients (n=12) (Mean). **C** Distribution of phenotypes P1-P7 subdivided in three groups based on Aβ proximity. Comparison of the prevalence of different clusters based on proximity to Aβ-plaques for cluster P1-P7 (**D-J** respectively). Example of infiltrating microglia of P4 (**K**) and P6 (**L**). For **B** and **D-J**, Stacked bars represent mean and statistical difference between spatial groups was tested using Repeated Measures ANOVA with Geisser-Greenhouse correction, individually for controls (n=9) and AD patients (n=12). *P*-values were corrected for multiple testing using Bonferroni for all test performed in B, and jointly on all test performed in D-J. Scale bars, 100 µm. Scale bar zooms, 50µm.

### 3.3 Morphology

Even though a large variety of activation-dependent microglia markers now exist, microglial morphology can add additional information on microglial activation. Therefore we evaluated the relationship between expression of the P2RY12, TMEM119 and Iba1 with the classically defined morphological states: homeostatic, activated (which also encompasses phagocytic and rod-shaped), and dystrophic microglia (19). Morphological evaluation was performed by visually assessing 5-10 microglia per image, in a total of 130 images (3-8 images per subject), and comparing the morphologically evaluated microglia with P2RY12, TMEM119 and Iba1 expression.

Iba1-expression was identified on all three microglia morphological subtypes. However, noticeable is that even though the majority of homeostatic microglia showed positive Iba1 signal, the expression of Iba1 on activated microglia was generally higher (Fig 4B vs. 4A). Dystrophic microglia only had higher Iba1 expression when there were signs of early dystrophy with spheroid formation, but enlarged soma were still visible. Once fragmentation of the branches occurred, usually Iba1 expression also was decreased (Fig. 4C-1). Finally, perivascular macrophages also showed consistent high Iba1 expression.

**Fig 4.**
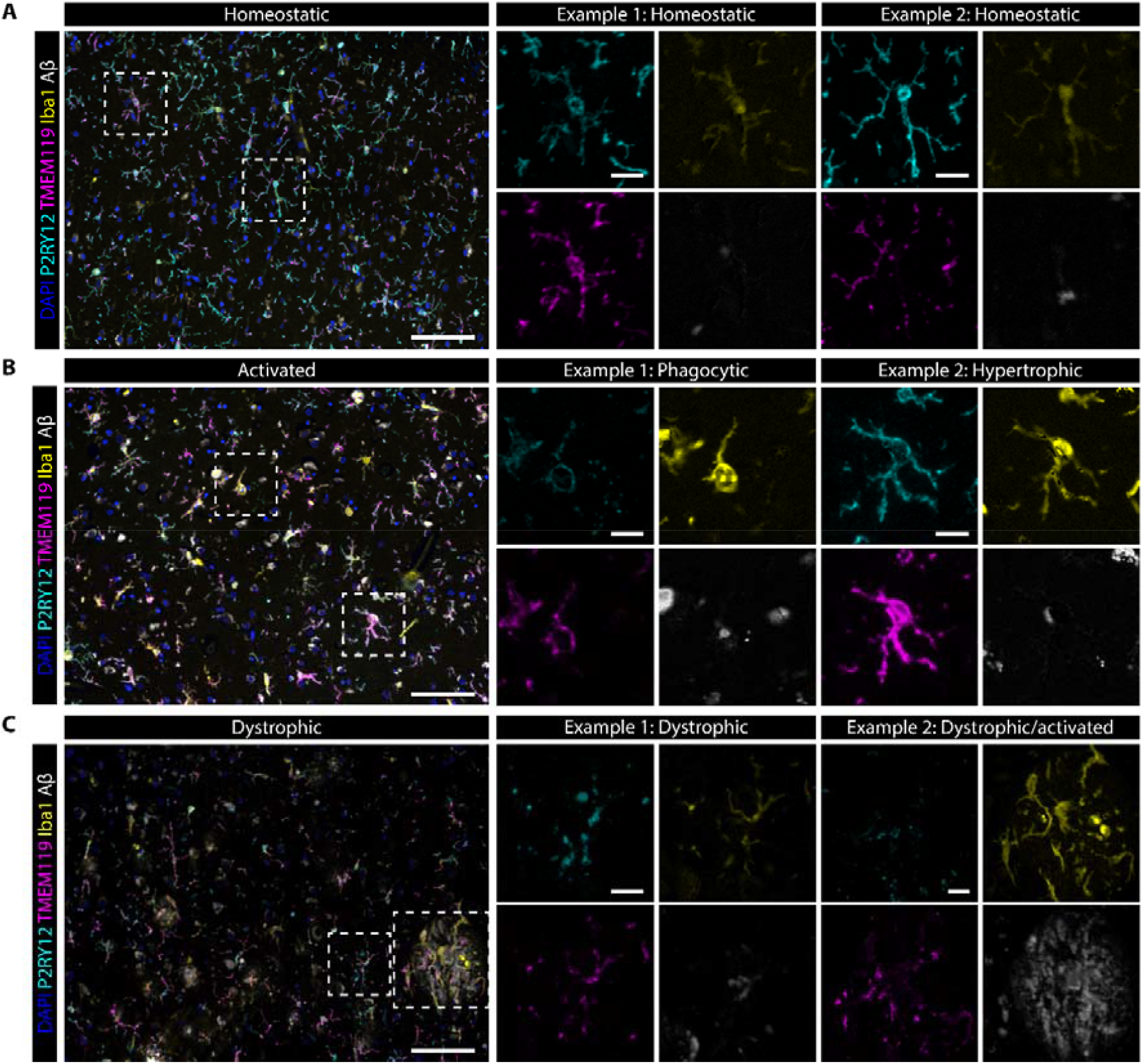
Correlation of marker-expression with morphology. Representative images showing a general appearance of homeostatic (**A**), activated (**B**) and dystrophic (**C**) microglia, with zooms showing individual marker-expression of microglia of a specific subtype. Scale bars, 100 µm. Scale bar zooms, 20 µm.

Strong P2RY12 expression was consistently found on homeostatic microglia (Fig. 4A). Also both activated and dystrophic microglia generally still expressed P2RY12, though not as consistently as homeostatic (Fig. 4B-1). However, a clear exception to this consistent expression were microglia surrounding Aβ-plaques (Fig. 4C), as described in the previous section. Apart from the spatial localization close to plaques, we did not observe clear morphological features associated with this specific plaque-related loss of P2RY12-expression.

TMEM119 was expressed on nearly all homeostatic microglia. Loss of TMEM119 was clearly found on a proportion of activated microglia, but not consistently across all (Fig. 4B-1 vs 4B-2). With regard to dystrophic microglia, a clear change in intracellular expression patterns was noticeable, with more punctate and irregular expression along the membrane, compared to homeostatic microglia. The clearest example of this disturbed distribution of TMEM119 was found in the Aβ-plaque infiltrating microglia (Fig. 4C-2).

All in all, none of the markers showed a consistent change in expression by morphological subtype of microglia. TMEM119 and P2RY12 were both consistently found on homeostatic microglia, but showed variable expression on activated and dystrophic microglia, although no unique morphological features correlated with expression levels of any of the markers. These results indicate that although morphology can clearly depict whether cells have lost their homeostatic function, the morphological subtypes do not accurately reflect specific activation states resulting in loss or gain of any of the markers.

## 4. Discussion

The present study is the first to quantify the co-expression patterns microglia specific markers TMEM119 and P2RY12, together with Iba1in human control and AD brains. We found unique patterns for each of the markers, with a selective decrease of TMEM119 and P2RY12, and an increase of Iba1 in subsets in AD patients. Spatially, distinct phenotypes preferentially infiltrated Aβ-plaques, and showed consistent Iba1 expression but marked loss of P2RY12, or both P2RY12 and TMEM119.

For Iba1, we found comparable numbers of positive cells in AD patients and controls, and it was consistently expressed in microglia surrounding and infiltrating Aβ-plaques. This was in line with a systematic review, showing no significant increase in the number of identified Iba1-positive microglia in AD patients in the majority of studies (17). However, when using expression-based assays (IHC/western blot/qPCR) some studies did show a slight increase of Iba1-expression in microglia in AD (17,20). In our study, by visually evaluating different morphological subtypes, we found that activated microglia appeared to have higher expression of Iba1, though quantitative evaluation of all raw expression values showed no differences between controls and AD patients. Additionally, upon further stratification of the microglia based on Aβ-plaque proximity, we did find a significant increase of the prevalence of Iba1 in Aβ-plaque surrounding and infiltrating microglia. Functionally, Iba1 is involved in the reorganization of the cell-skeleton by cross-linking actin to induce membrane ruffling (21), essential for both microglia motility and phagocytosis (22). These are two principal features of activated microglia, potentially explaining the observed increase of Iba1 in the morphologically activated and Aβ-plaque associated microglia in our study. Via co-staining for multiple markers we could verify that specific Iba1^+^-microglia phenotypes are variably increased and decreased in AD, but do not result in an overall significant difference in prevalence as, reported in the systematic review(17). Finally, even though Iba1 is often considered as a pan-marker for all microglia and macrophages (17,19,23,24), we observed a significant amount of microglia that were Iba1^-^ (10-60%), especially in our controls. This discrepancy could be explained by the fact that studies with single Iba1 IHC or double-staining of Iba1 with P2RY12 or TMEM119 would have missed microglia solely expressing P2RY12 or TMEM119 (P1-2), resulting in a higher percentage of Iba1^+^ microglia. This had already been suggested by Hendricks *et al*, who qualitatively showed only scarce expression of Iba1 in ramified microglia in control tissue, and Iba1 not being expressed by all microglial activation stages (25). Two other quantitative IHC studies using double stainings also described populations of microglia showing low Iba1 expression (26,27). However, contrary to our results and that of Hendricks *et al*, they found these Iba1^low^ populations to be increased in AD. All in all, we can confirm quantitatively that Iba1 is not expressed on all microglia, though opposing literature exists on whether activated microglia shows increased or decreased expression.

Regarding P2RY12, in our study we did not find significant differences in the overall number of P2RY12^+^-microglia. Nevertheless, specific phenotypes P2RY12^+^ phenotypes (P1 & P5) were found to be more prevalent, whereas others (P7) were less prevalent in AD. Correspondingly, a previous study by Swanson *et al*, showing the first quantification of P2RY12 expression in AD brains using single-cell histology analysis, also found no difference in the prevalence of microglia populations with high or low P2RY12 expression between controls and AD patients (27). Further spatial investigation showed that P2RY12^+^-phenotypes were specifically decreased in and surrounding Aβ-plaques (P1 & P3), where P2RY12^-^-phenotypes were more prevalent (P4, P6), which contributed to an overall loss of P2RY12 microglia infiltrating Aβ-plaques. These findings confirm previous observations of areas lacking P2RY12^+^-microglia around Aβ-plaques in AD patients (15,16). Functionally, it was found that the P2Y12-receptor can induce microglial chemotaxis after binding of ligands such as ATP/ADP or extracellular nucleotides (28,29). This indicates P2RY12 to play an important role during early stages of microglia activation in response to CNS injury or blood-brain barrier damage-damage (29,30), after which expression is lost. *In vitro* results are in line with this, showing P2RY12 downregulation after activation with LPS (31,32). Also for multiple sclerosis this pattern is observed, with general decrease of P2RY12 expression, and almost complete loss in white matter lesions (15,24,31). Interestingly, in our human post-mortem AD tissue, P2RY12 was still variably expressed throughout the brain by microglia with both activated or dystrophic morphological appearance, but only specifically lost surrounding Aβ-plaques. From this one can conclude that although P2RY12 appears to be a protein predominantly expressed in homeostatic microglia, its expression is only lost under very specific conditions, and presence of P2RY12 does not conclusively indicate a homeostatic function.

Lastly, a significant reduction of TMEM119^+^-microglia and expression levels was found in AD. Interestingly, in contrast to P2RY12, this was generally not associated with Aβ-plaques. Though one subset of microglia which was significantly associated with Aβ-plaques showed loss of both TMEM119 and P2RY12-expression (P5), another Aβ-associated subset still expressed TMEM119 (P7). Notably, even though TMEM119 was still expressed in this subset of microglia, the protein appeared to be redistributed and did not show a homogenous appearance along the membrane as observed in controls patients. To date, only few have studied the expression of TMEM119 in microglia in aged and AD patients. Firstly, in line with our results, Swanson *et al* found a significant increase of a microglia population with low TMEM119 expression in AD patients. Secondly, using RT-PCR, an increase of TMEM119 mRNA was observed in AD brains, though this could not be confirmed on western blot (13). In other diseases such a multiple sclerosis, researchers found a significant decrease in white matter lesions (13,31), but a different study interpreted this as the cells being macrophages (24). Additionally, very little is known about the function of TMEM119 in microglia. Consequently, TMEM119 is still occasionally incorrectly considered to be consistently expressed by microglia under both homeostatic and disease conditions, and used to discriminate microglia from blood-derived macrophages. However, our results indicate that TMEM119 expression is also dependent on activation state; its expression is decreased in an activated, though partially disparate subset than P2RY12^-^-microglia. Cells lacking TMEM119 expression but with clear Iba1 and or P2RY12 expression were consistently observed in the parenchyma and showed clear microglia morphology. This indicates that a lack of expression of TMEM119 on an Iba1-positive cell does not necessarily indicate that the observed cell is not of microglial origin.

Finally, this works helps interpret studies previous and future studies using Iba1, TMEM119 and P2RY12 or a combination of these markers. We have therefore created a figure (Fig. 5) to briefly summarize our findings and help interpret one’s own study. The occurrence of different combinations of markers appeared heterogeneous, likely reflecting the myriad of functions microglia execute, and not general morphological activation states. Additionally, based on this study one can conclude that when evaluating microglia numbers, it is important to always stain for multiple different microglia markers, as none of the markers are expressed by all microglia, and an observed difference in microglia numbers could be due to a loss of marker expression, rather than loss of microglia. The same applies when studying macrophage-infiltration as lack of TMEM119 does not immediately indicate macrophage rather than microglial origin.

**Fig. 5.**
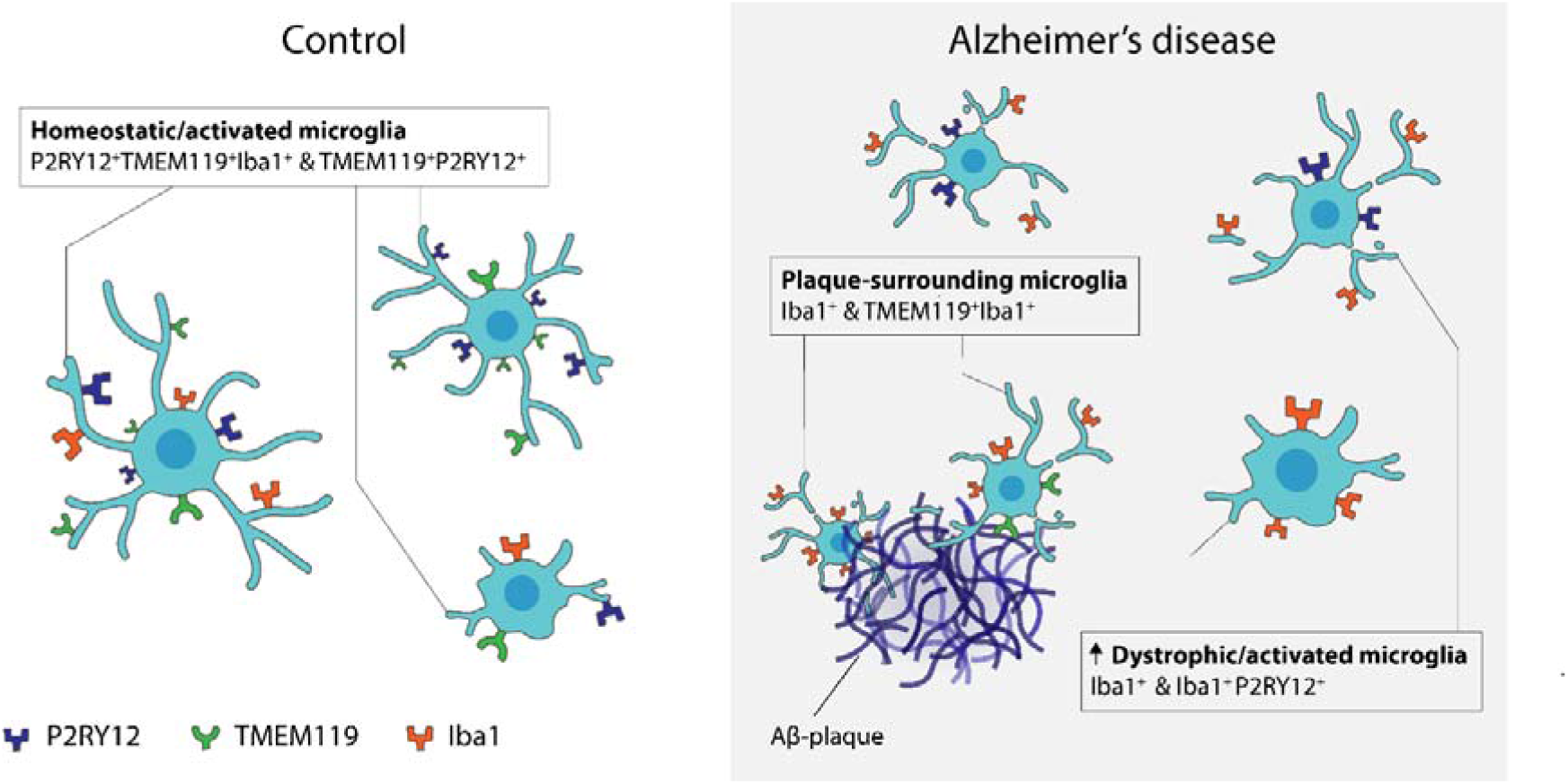
Schematic of observed findings in control and AD patients. In controls the majority of microglia express all three markers or TMEM119 and P2RY12. On the other hand, microglia solely expressing Iba1 are more present in AD patients, as are microglia expressing Iba1 with either TMEM119 or P2RY12. These results indicate a shift from microglia with predominant TMEM119/P2RY12 expression to a microglia state with more Iba1 expression and loss of TMEM119 and/or P2RY12 in AD patients. Loss of P2RY12 was specifically attributed to presence of Aβ-plaques, whereas TMEM119 was more generally lost in both grey and white matter.

All in all, this works aids to the body of literature available on immunohistochemical markers used for microglia research. By employing multispectral immunofluorescence to specifically study the co-expression patterns of these microglia markers enabled us to show the overlap and differences in expression in control aged and diseased brains, and in relation to Aβ-plaques. Thanks to the automated microglia identification and marker evaluation, this work could easily be extended in the future with more recently identified microglia markers such as Hexb and Siglec-H (33,34).

## Abbreviations

Aβ: Amyloid-β
AD: Alzheimer’s disease
CNS: Central Nervous System
Iba1: Ionized calcium binding adaptor molecule 1
IHC: Immunohistochemistry
TMEM119: Transmembrane protein 119

## Ethics approval and consent to participate

All material has been collected with written consent from the donors and the procedures have been approved by the Medical Ethical committee of the LUMC and the Amsterdam UMC.

## Consent for publication

Not applicable

## Acknowledgements

We acknowledge all patients who donated their brain to the Leiden University Medical Center (LUMC), Netherlands Brain Bank (NBB) or the Normal Aging Brain collection Amsterdam (NABCA), and prof. A.J.M. Rozemuller for neuropathological evaluation of the brains. We would like to thank the department of Pathology of the LUMC, especially Marieke IJsselsteijn and Noel de Miranda for their help with setting up multispectral immunofluorescence for microglia. We owe thanks to O. Dzyubachyk, B.P.F. Lelieveldt and J. Dijkstra for support in the analysis of the multispectral immunofluorescence data.

## Data availability

Data sharing is not applicable to this article as no datasets were generated. Scripts for data analysis are available from the corresponding author upon reasonable request.

## Authors’ contributions

B.K. and L.v.d.W. conceived and designed the project. B.K. acquired the multispectral immunofluorescence (mIF) data. A.S. and B.K. created the microglia segmentation pipeline. B.K. and L.R.T.K. assessed data-quality and manually scored subsets of data. A.S. created the spatial analysis tools for mIF data under supervision of T.H.. B.K performed morphological evaluation of microglia. B.K. and A.S. analysed and interpreted the mIF data. B.K., W.M.C.v.R-M, and L.v.d.W. wrote the manuscript. All authors read and approved the final manuscript.

