## Supplementary file for "Co-expression patterns of microglia markers Iba1, TMEM119 and P2RY12 in Alzheimer’s disease"

### Supplementary files

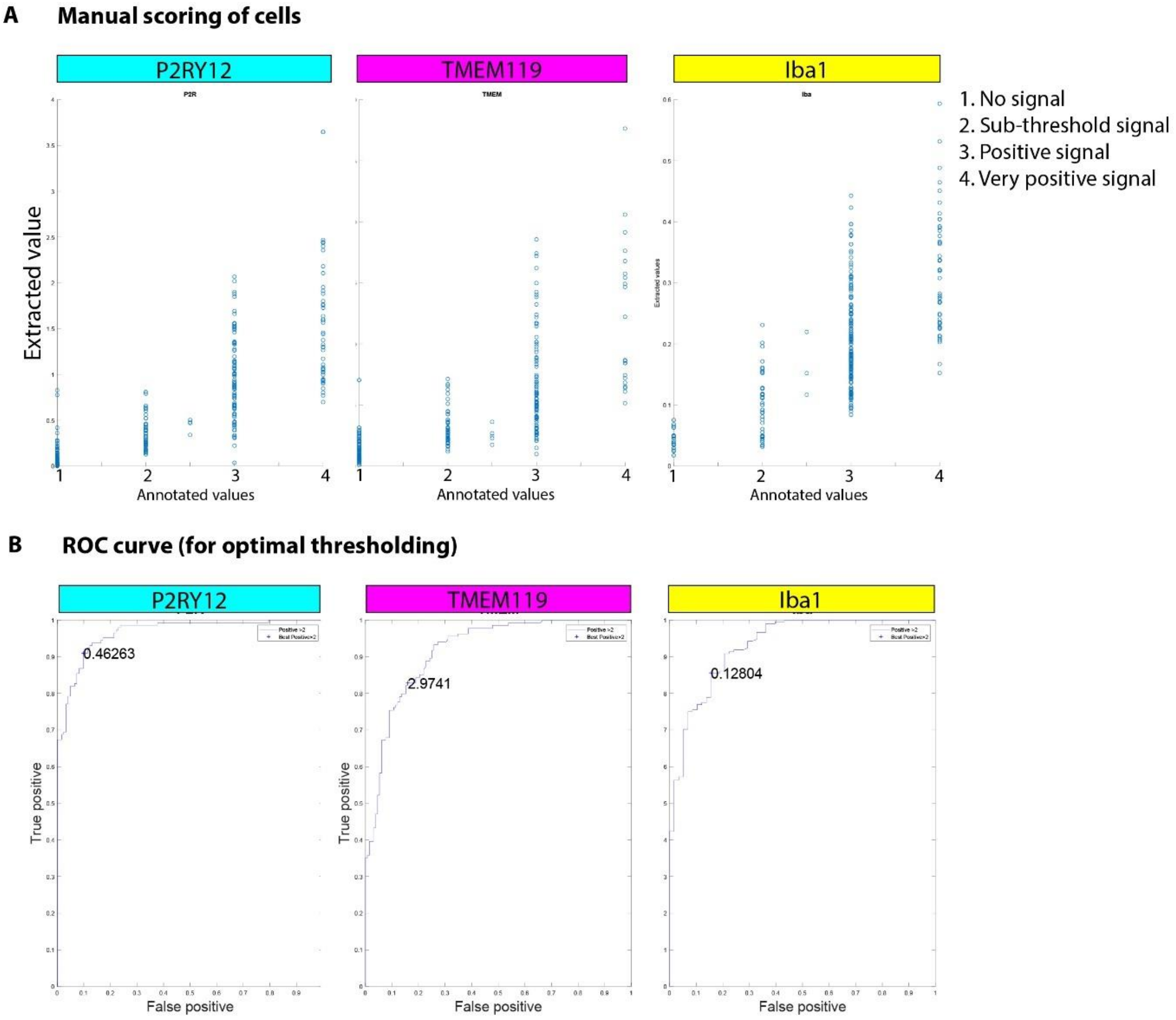


**Supplementary Fig. 1 Manual thresholding of cells. A** Annotated values of each individual cell plotted against the extracted raw fluorescent intensity value. **B** ROC curve showing true positive and false positive rate for a thresholding-value to distinguish between positive and negative cells based on the manual annotation.

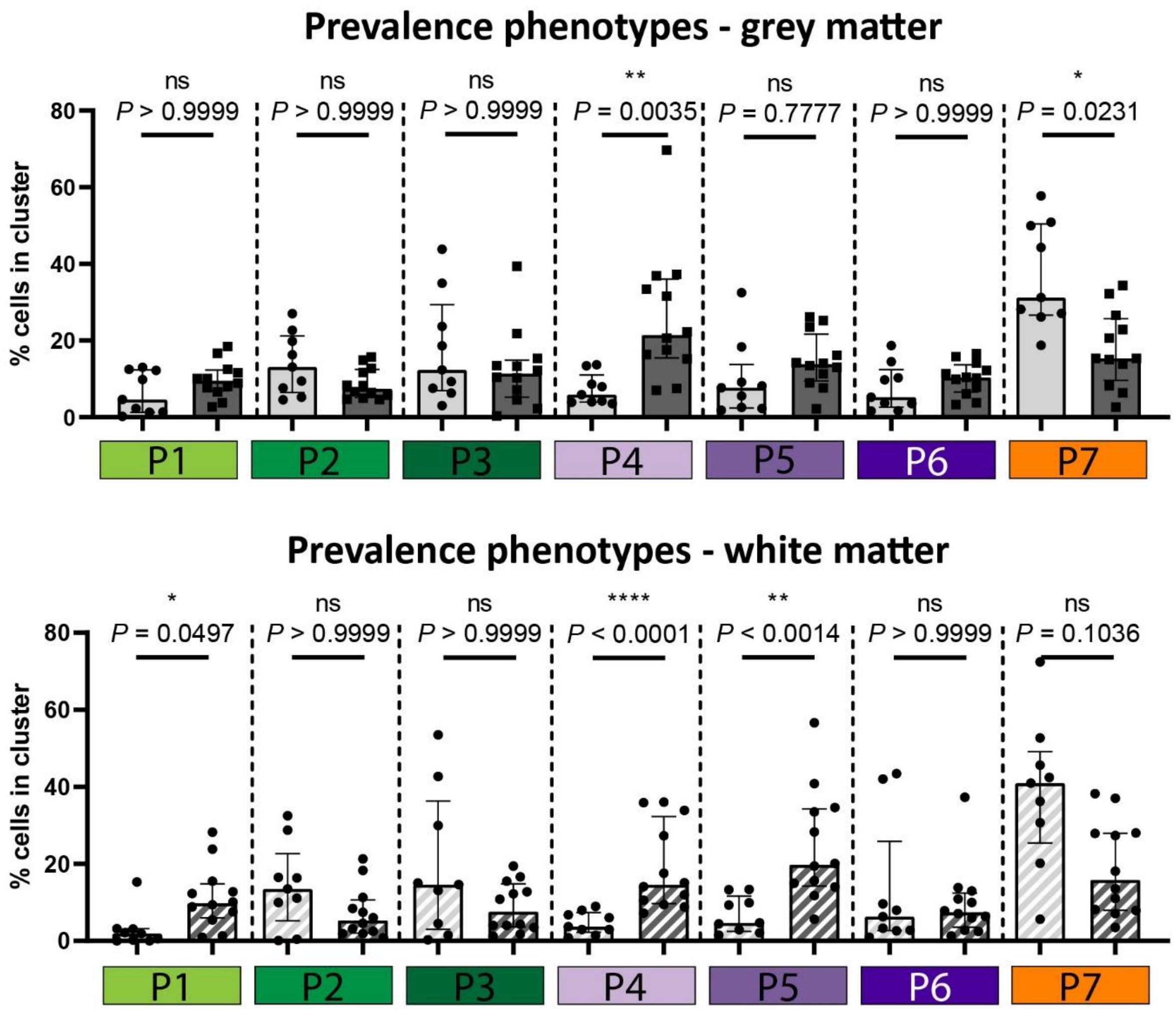


**Supplementary Fig. 2 Prevalence of microglia phenotypes in grey- and white matter.**

Median, Mann-Whitney U test. *P*-values were corrected for multiple testing using Bonferroni for grey matter and white matter phenotypes separately.
